# Differences in working memory coding of biological motion attributed to oneself and others

**DOI:** 10.1101/2021.08.16.456121

**Authors:** Mateusz Woźniak, Timo Torsten Schmidt, Yuan-hao Wu, Felix Blankenburg, Jakob Hohwy

## Abstract

The question how the brain distinguishes between information about oneself and the rest of the world is of fundamental interest to both philosophy and neuroscience. This question can be approached empirically by investigating how associating stimuli with oneself leads to differences in neurocognitive processing. However, little is known about the brain network involved in forming such self-associations for, specifically, bodily stimuli. In this fMRI study, we sought to distinguish the neural substrates of representing a full-body movement as one’s movement and as someone else’s movement. Participants performed a delayed match-to-sample working memory task where a retained full-body movement (displayed using point-light walkers) was arbitrarily labelled as one’s own movement or as performed by someone else. By using arbitrary associations we aimed to address a limitation of previous studies, namely that our own movements are more familiar to us than movements of other people. A searchlight multivariate decoding analysis was used to test where information about types of movement and about self-association was coded. Movement specific activation patterns was found in a network of regions also involved in perceptual processing of movement stimuli, however not in early sensory regions. Information about whether a memorized movement was associated with the self or with another person was found to be coded by activity in the left middle frontal gyrus (MFG), left inferior frontal gyrus (IFG), bilateral supplementary motor area, and (at reduced threshold) in the left temporoparietal junction (TPJ). These areas are frequently reported as involved in action understanding (IFG, MFG) and domain-general self/other distinction (TPJ). Finally, in univariate analysis we found that selecting a self-associated movement for retention was related to increased activity in the ventral medial prefrontal cortex.

## Introduction

The body is considered as the basis of our self – as Baumeister (1999) influentially remarked: “Everywhere in the world, self starts with body”. While the abstract, language-mediated aspect of our self emerge only later in our lives, knowledge about our bodies is present from the onset of our existence. These two types of information have been argued to form two distinct self-representations: the bodily self and the conceptual self (Christoff, Cosmelli, Legrand, & Thompson, 2011; Gallagher, 2000; Neisser, 1988). This difference also finds support in research on the brain mechanisms underpinning processing of bodily and abstract self-related information. While the bodily self is underpinned primarily by multisensory and sensorimotor brain systems (Blanke, 2012; Blanke, Slater, & Serino, 2015; Christoff et al., 2011; Limanowski & Blankenburg, 2013; Tsakiris, 2010, 2017), the conceptual self is mostly related to the activity of higher-level brain areas, especially the cortical midline structures (Hu et al., 2016; Legrand & Ruby, 2009; Martinelli, Sperduti, & Piolino, 2013; Northoff et al., 2006; Qin & Northoff, 2011; Yin, Bi, Chen, & Egner, 2021), and there is relatively little overlap between the brain areas involved in representing these two types of self-related information (Hu et al., 2016; Qin, Wang, & Northoff, 2020).

Neurocognitive processing of information about one’s bodily self is typically investigated through the use of images or videos of participants’ face, body, body parts, or body movements. However, all of these stimuli are characterized not only by their self-relatedness, but also by the fact that they are highly familiar to participants. Most previous studies have attempted to minimize the influence of familiarity by comparing processing of self-related stimuli (own face, own movement pattern) with stimuli representing familiar or famous others (friends, family, celebrities). However, this kind of control conditions can never fully match with regard to familiarity. We spend all of our lives inside our bodies, while we devote substantially smaller fraction of lifetime paying attention to the bodies of other people, even famous ones. Therefore, while studies comparing processing of self-related information with information related to familiar others provide an important contribution to our understanding of how the brain represents our bodily self, there is a need for validation of these results using designs which to a greater extent control for familiarity.

To control for familiarity in self-related processing, recent studies have begun investigating how creating ad hoc self-associations can modulate cognitive processing. In such studies participants are told to associate arbitrary stimuli (e.g. geometrical shapes) with various identities. Crucially, one of these identities is the participant themself. For example, a participant can be asked to associate a triangle with themself (through an instruction: “You are a triangle”), a square with a friend, and a circle with a stranger (Sui, He, & Humphreys, 2012; Sui, Rotshtein, & Humphreys, 2013). The results show that even an arbitrary association with the self leads to robust prioritization of self-associated stimuli, as manifested by shorter reaction times and improved detection accuracy. In behavioral studies, the same procedure has been also used with bodily stimuli such as faces and avatar bodies, yielding similar effects (Mattan, Quinn, Apperly, Sui, & Rotshtein, 2015; Payne, Tsakiris, & Maister, 2017; Woźniak & Hohwy, 2020; Woźniak & Knoblich, 2019). At the neural level, several previous experiments investigated which brain areas are affected by creating such arbitrary self-associations for abstract, symbolic stimuli (Lockwood et al., 2018; Sui, Liu, Mevorach, & Humphreys, 2015; Sui et al., 2013; Yankouskaya et al., 2017; Yankouskaya & Sui, 2021; Yin et al., 2021; Zhao, Uono, Li, Yoshimura, & Toichi, 2018) and found that midline cortical structures (especially the vebtromedial prefrontal cortex) are critically involved in such tasks. However, to date, no neuroimaging study used this approach to investigate which brain areas represent bodily information associated with the self. This is a significant omission, as bodily self is arguably more basic than the conceptual self (Christoff et al., 2011; Gallagher, 2000; Gillihan & Farah, 2005), and also because, as mentioned before, previous research found that brain regions underpinning aspects of one’s bodily self rarely overlap with brain regions responsible for processing of abstract self-related information (Gillihan & Farah, 2005; Hu et al., 2016; Qin et al., 2020).

The present study investigated, focusing on full-body movements, which brain regions differentially represent self-versus other-associated bodily information. Specifically, it was designed to address how the brain differentiates between seeing identical actions – i.e. the same movements presented from a third-person perspective – attributed to oneself or to other people. Participants were presented with stimuli depicting different types of movements, presented as point-light walkers (short animations presenting people performing an action, but devoid of any information about their physical body appearance). Critically, across different trials participants were told to attribute the same movements either to themselves, or to others, which serves to match familiarity between the conditions. We employed a working memory paradigm (delayed-match-to-sample task) during which participants had to form and actively maintain the representation of a self or non-self-associated movement for several seconds. We used MVPA on fMRI data to detect brain areas that differently code memorized movement associated with the self or another person.

Previous research on self-related processing suggest that it is mainly activity in three brain regions distinguishing whether a presented movement is associated with oneself or other identities. First, selective processing of self-related information have been reported in frontal cortices that are also involved in action recognition and which are believed to be part of the human mirror-neuron system, especially the inferior frontal gyrus (Molenberghs, Cunnington, & Mattingley, 2012). Second, the ventral medial prefrontal cortex has been found to play an important role in processing abstract self-related information, such as one’s name, self-related adjectives, or arbitrary self-associated stimuli (Hu et al., 2016; Sui et al., 2013). Finally, the temporal parietal junction and surrounding areas have been identified as being critically involved in self-other distinction across a range of cognitive processes (Ganesh, van Schie, de Lange, Thompson, & Wigboldus, 2011; Hu et al., 2016; Quesque & Brass, 2019; Sui et al., 2013). Based on these reports, we were particularly interested in the involvment of these three regions in representing movements as self-related. To test this, we included three experimental conditions, namely memorizing the movement of oneself, or of two different stranger identities. This design allowed us to test for distinguishable activation patterns evoked by the self-associated and stranger-associated movements, while we expected no differences between the two strangers.

## Methods

### Participants

Twenty-four volunteers participated in the study (age range: 21 - 40 years; *M*=26.3, *SD*=4.89; 13 females, 11 males). All participants were right-handed. All participants had normal or corrected-to-normal vision. Informed consent was obtained from all participants before the start of the experiment according to procedures approved by the Freie Universität Berlin Ethics Committee.

### Procedure

The experimental paradigm comprised a delayed match-to-sample working memory task employing retro-cues associated with specific identities. Inside the MRI scanner, participants first practiced the task while structural MRI data was acquired. They trained until they felt confident and showed a task performance of at least 70% correct in a 5 minutes long practice block (if they failed to reach this threshold they had to repeat the practice). Next, five experimental runs of 15 min 30 seconds each were conducted while functional MRI data were collected.

At the beginning of the experiment, participants learned which of three colors (red, green, blue) represents which one of three identities: You (the participant her- or him-self), Pat (Patricia or Patrick – gender of the name was always matching with participant’s gender), and Doc (a gender-matched stranger who is a doctor). The labels “Pat” and “Doc” were chosen as indicators of two strangers’ identities because they have the same length as the word “You”, they can be used to reflect both genders (so stranger-identities were always of the same gender as the participant), and each belongs to a different grammatical category ruling out potential confound effect of grammatical distinctiveness (Schäfer, Wentura, & Frings, 2017). The associations between colors and identities were presented in a written form on the instruction sheet which was given to each participant. The color-identity associations were counterbalanced across participants using a Latin square scheme. Color-identity associations were introduced to the task in order to avoid using labels as retro-cues in the experimental task.

Figure 1 illustrates the procedure of the retro-cue task used in the present study. In each trial participants had to memorize a biological motion stimulus, which was indicated to be either a movement of oneself or of another person. Each trial started with the presentation of two short animations of movements displayed as point-light walkers (each 1000 ms, separated by a 1000 ms ISI, see section Stimuli). Before each stimulus the identity performing the motion was indicated by an identity-cue: either “YOU” (self-condition), “PAT”, or “DOC” (others-conditions) for 500ms (see **Figure 1**). For example, if the first cue was “PAT” then it indicated that the first movement was performed by Pat, and if the second cue was “Doc” then it conveyed the information that the second movement was performed by Doc. If one of the cues was “You” then participants were told to interpret the corresponding movement as if they were watching themselves from a third-person perspective (for example on a video recording). After the presentation of stimuli, a retro-cue instructed which of these two movements should be maintained for the working memory phase. The retro-cue was either a red, blue, or green circle and indicated the identity (You, Pat, or Doc) whose movement had to be memorized, according to the previously learned color-identity mappings. For example, if the associations were the same as presented in **Figure 1A** (upper right) then a blue retro-cue told a participant to memorize Doc’s movement, and if it was red then they were expected to memorize “Your” (participant’s) movement. After a 7 second retention period, a target movement was presented and participants’ task was to decide if the target movement was identical (i.e. the same type of movement and presented from the same point of view) to the memorized movement. Reports were given via right-hand middle and index finger button presses (yes/no-left/right balanced across participants).

**Fig 1.**
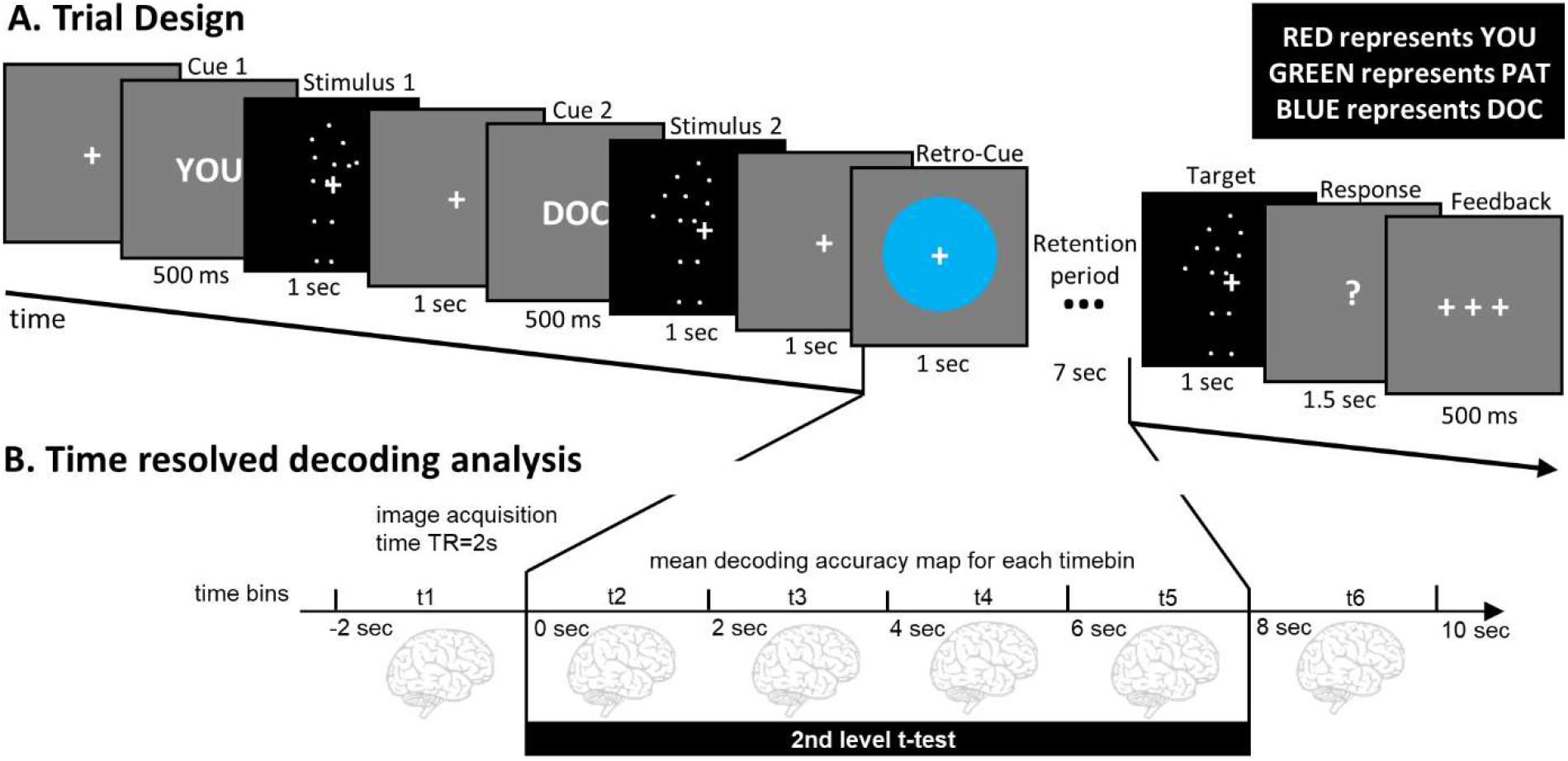
Experimental paradigm and analysis. **A**. During fMRI scanning, participants performed a retro-cue delayed match-to-sample working memory task. In each trial, two full-body movement stimuli were presented and a consecutive retro-cue indicated which of the two movements had to be retained for a 7 sec retention period. Finally, a target movement stimulus was presented and had to be judged whether it was the same or different from the memorized movement. A distinctive feature of this task was that each of two presented movements was associated with one of three identities: either with the participant oneself or with one of two strangers. The color retro-cue indicated which movement should be memorized by referring to the identity of a person performing a movement. Assignment of colors (example given in upper right panel) was presented to participants in the instructions to the task and was randomized across participants. **B**. As fMRI data acquisition was time-locked to the start of the working memory retention period, it was possible to conduct time-resolved decoding analysis. In short, whole-brain searchlight classification analyses were performed on the data of each timebin, corresponding to separate functional volumes. We used group-level t-contrast across timebins t2-t5 to test for above-chance decoding accuracies.

Colors rather than labels were used as retro-cues to avoid potential confounds related to label-specific processing, as there exists evidence that self-related labels are processed preferentially over other labels (Hu et al., 2016; Shi, Zhou, Han, & Liu, 2011; Woźniak & Knoblich, 2019; Woźniak, Kourtis, & Knoblich, 2018; Zhou et al., 2010). A target movement was only considered identical to the memorized movement if it was both the same type of movement (e.g. chopping, painting) and was presented from the same camera position, that is, it was identical in regard to low-level visual features. Participants were required to make their responses within a time window of 1.5 s after the target movement offsets. Afterwards they were presented with feedback provided as “+ + +” (correct), “- - -” (incorrect), or “0 0 0” (missing response). The inter-trial interval was either 1s or 3s, balanced and randomly interleaved across trials.

Each run comprised 48 trials which were balanced with regards to: a) which movement was retained in working memory, b) whether the first or the second movement was retained, and c) which identities were presented as cue 1 and cue 2. The target movement was identical to the memorized movement in 50% of the trials, a different type of movement in 25% (randomly selected) of the trials and in the remaining 25% it was the same type of movement, but presented from a different camera position. The remaining factors were counterbalanced across trials (which movement was presented as a target, which camera position was used for each movement, etc.).

### Stimuli

The experiment was programmed and conducted using Matlab (version R2013 8.2). The Psychophysics Toolbox (version 3.0.10) was used for presentation of the experimental task, and Cogent 2000 Toolbox (developed by the Cogent 2000 team at the FIL and the ICN) was used to create log files.

Short movie clips presenting point-light walkers illustrating eight different movements were chosen from the data set created by (Vanrie & Verfaillie, 2004). All stimuli portrayed full bodies performing movements with their hands while standing (chop, drink, mow, paint, peddle, spade, stir, sweep) from five different camera positions (frontal, 45° left, 90° left, 45° right, 90° right). Because original clips differed in length, only one second (30 frames) was selected from each of them. For each participant, four of these eight movement stimuli were selected as the stimulus set. The assignment of movements to participants was pseudo-random in a way that ensured that each movement type was used for an equal number of participants.

Visual stimuli were projected with an LCD projector (800 x 600, 60 Hz frame rate) onto a screen on the MR scanner’s bore opening. Participants observed the visual stimuli via a mirror attached to the MR head coil from a distance of 110 ± 2 cm. Fixation cross and visual cues were presented in white on a grey background. All identity-cues were presented in font size 14; the radius of the retro-cue was 10 px. The size of point-light walker stimuli was 150×200 px.

### Data acquisition and preprocessing

MRI data were acquired using a 3T TIM Trio scanner (Siemens, Erlangen) at the Center of Cognitive Neuroscience Berlin at Freie Universität Berlin. A structural T1-weighted MRI image was acquired (MPRAGE; 176 sagittal slices/3D acquisition; TR = 1900ms; TE = 2.52ms; 1×1×1 mm^3^ voxel; flip angle = 90°; field of view = 256mm) when participants were practicing the task during the practice run. Functional MRI data were acquired in five runs, each lasting 15 minutes 30 seconds. In each run, 565 images were acquired (T2*-weighted gradient-echo EPI; 37 slices; ascending order; 20% gap; whole brain; TR=2000ms; TE = 30ms; 3×3×3 mm^3^ voxel; flip angle = 70°; field of view = 192mm; 64×64 matrix). Trial onsets as well as onsets of retro-cues and retention periods were time-locked to the onset of functional image acquisition allowing TR-wise analysis (See **Figure 1**).

### Data analysis

Preprocessing of the data and GLM analyses were conducted using SPM12 (Wellcome Trust Centre for Neuroimaging, Institute for Neurology, University College London, London, UK). Preprocessing of functional data for decoding analyses was limited to spatial realignment and temporal detrending (Macey, Macey, Kumar, & Harper, 2004), to preserve the spatiotemporal structure of data for the decoding analysis.

#### Time-resolved searchlight decoding

We used a time-resolved multivariate searchlight decoding approach as applied in previous working memory studies (Christophel, Hebart, & Haynes, 2012; Kriegeskorte, Goebel, & Bandettini, 2006; Schmidt, Wu, & Blankenburg, 2017). We performed two independent analyses on the data, in order to determine brain areas that (1) carry information about the memorized movement as such (Movement Decoding: MD) and (2) the identity of the person performing a memorized movement (Identity Decoding: ID).

For this purpose we first fitted two independent subject-level GLMs comprised of finite-impulse-response (FIR) regressors (high-pass filter cut-off: 200 sec) modelling the time period from 2 sec before the onset of the retro-cue to 2 sec after the WM retention period, corresponding to 6 FIR regressors of 2 sec each (See **Figure 1B**). The first GLM comprised regressors for the three identities (You, Pat, or Doc). The second GLM modelled the retention activity with regressors for the different movements irrespective of the identity. This was possible, as the factors movement type and identity were orthogonal in our experimental design. The FIR GLM conducted for Identity Decoding resulted in 95 beta estimates (3 conditions × 6 time bins × 5 runs + 5 constants), and the FIR GLM for Movement Decoding yielded 125 beta estimates (4 stimuli × 6 time bins × 5 runs + 5 constants). Beta estimates were forwarded to the subsequent multivariate pattern analysis.

We used The Decoding Toolbox (TDT) (Hebart, Görgen, & Haynes, 2015), which provides an interface for applying machine learning algorithms to neuroimaging data using LIBSVM library (Chang & Lin, 2011). We combined MVPA with a searchlight routine (Kriegeskorte et al., 2006), which allowed us to conduct data-driven analysis without any prior assumptions on the localization of information. The spatially-unbiased searchlight MVPA were performed for each of the six time bins. In the searchlight analysis, a spherical region-of-interest (radius = 4 voxels) was moved voxel-by-voxel through the measured brain volume. At each position, we used Support Vector Machine (SVM) to test for the decodability of WM contents in a pair-wise manner (linear kernel with constant regularization parameter c=1). Beta estimates for each time bin were z-scaled (normalized) across the samples for each voxel as implemented in TDT. The decoding analyses were performed using a five-fold, leave-one-run-out cross-validation decoding scheme across the five experimental runs. This procedure yields whole-brain decoding accuracy maps for each subject and each time bin. Decoding accuracy maps were normalized to MNI space, using unified segmentation and spatially smoothed using a 5mm FWHM Gaussian kernel, using SPM 12 routines. For the Movement Decoding analysis accuracy maps for each comparison between four movements (6 comparisons per participant) were averaged to obtain one decoding accuracy map per timebin. For the Identity Decoding analysis accuracy maps for comparisons between the participant and other identities (You vs. Pat and You vs. Doc) were averaged to obtain one map of You vs. Others comparison per timebin. The accuracy maps for comparison Pat vs. Doc for each timebin were used as an independent control condition contrasting two stranger identities.

We obtained six accuracy maps for Movement Decoding (reflecting decoding in six time bins), and twelve accuracy maps for Identity Decoding: six maps reflecting decoding of self vs. others (Pat and Doc), and six maps reflecting decoding of Pat vs Doc. The resulting accuracy maps were entered to a SPM flexible-factorial repeated-measures second-level design (separately for Movement Decoding and Identity Decoding) to test for above chance decoding accuracies across the six timebins. Consequently, we ran three independent group-level ANOVAs: (1) Movement Decoding, (2) Identity Decoding: self vs. others, and (3) Identity Decoding: Pat vs. Doc. We calculated t-contrasts against chance decoding accuracy (50%) across timebins 2, 3, 4, and 5 (see Figure 1). Results are reported at *p* < 0.05, family-wise error (FWE) corrected for multiple comparisons, with a minimal cluster extent threshold of 50 voxels.

### Control analysis: decoding of non-memorized stimulus

In order to test for the specificity of our main decoding analyses, we conducted control analyses to test for above chance decoding of the stimulus which was not retained for working memory (uncued stimulus). New FIR models were estimated which, due to the balanced experimental design, had the same amount of regressors, as each stimulus was equally often memorized as they were presented and not memorized. We performed analogous decoding analyses for the non-memorized (1) Movement Decoding, (2) Identity Decoding: self vs. others, and (3) Identity Decoding: Pat vs. Doc.

#### Univariate analysis

A univariate analysis was conducted in order to identify regions that exhibit increased activity when processing self-than other-related movements. The preprocessed data were normalized to the MNI space and spatially smoothed using a 5mm FWHM Gaussian kernel, using SPM 12 routines. High-pass filtered data (cut-off: 128 s) were used for the model. On the subject-level, we modelled ten regressors. We included one regressor modeling the influence of all cues and movements presented before the retro-cue, three regressors for the onset of presentation of a retro-cue in each trial, independently for self-, Pat- and Doc-associated working memory content, three regressors for the period of working-memory retention, also independently for self, Pat and Doc, and finally one for the onset of the target, and two to model participants’ responses (independently for left- and right-hand responses). Each of these regressors was convolved with a canonical haemodynamic response function which took into account the duration of each type of regressor (e.g 1 second for a retrocue, 7 seconds for the retention period). Consecutively, a random-effects group-level analyses were conducted to calculate t-contrasts between self- and others-associated activity a) elicited by the retro-cue, and b) elicited during the working memory retention activity, using relevant subject-level contrasts (self vs. others). An analogous control group-level analyses were conducted for Pat vs. Doc comparisons. Similarly to decoding analyses, we report only clusters extending over at least 50 voxels with p<0.05 FWE.

## Results

### Behavioral performance

Participants performed with a mean of 83.6% (S*D*=5.7%) correct responses in the applied match-to-sample task (range: 70% - 94.2%). Testing for differences in performance levels (percentage of correct responses) with a one-way repeated measured ANOVA with factor identity (self, Pat, Doc) revealed a significant main effect of identity (*F*(2, 46)=5.49, *p*=0.007, partial η^2^=0.19). Post-hoc t-tests showed that participants were significantly less accurate to recognize a self-associated movement (*M*=81.7%, *SD*=5.4%) than a movement associated with Pat (*M*=84.5%, *SD*=6.3%, t(23)=3.38, p=0.008 Bonferroni-corrected) or Doc (*M*=84.4%, *SD*=7.2% t(23)=2.68, p=0.04 Bonferroni-corrected), while the difference between Pat and Doc was not significant (t(23)=0.1, *p*=1 Bonferroni-corrected).

In order to compare accuracies for different types of movements we conducted a linear mixed model (LMM) analysis. Conducting an LMM analysis was necessary in order to account for the crossed-nested character of this data. The analysis did not reveal any significant differences between movements. We conducted an LMM analysis using Statsmodels Python API with dependent variable being accuracy (percentage of correct responses), each of eight movements used in the experiment as independent variables, and participant as a grouping factor. We did not find any significant differences between accuracies for memorizing different types of movements (all movements were compared against the movement with highest accuracy, i.e. chopping).

### Movement Decoding

To identify regions where brain activation relates to the movement presented in the stimulus, we computed a t-contrast to test for above-chance decoding accuracies during the WM retention. The results (at p<0.05 FWE, min. cluster size = 50) are displayed in Table 1 and **Figure 2**. The largest cluster encompasses a big part of the parietal cortex, as well as occipital and temporal cortices (all bilaterally), with peaks in the left (posterior) middle temporal gyrus, left superior parietal lobe, and right inferior occipital gyrus. The second-largest cluster spans the bilateral precentral gyrus as well as the supplementary motor area.

**Table 1.** Regions with above-chance decoding accuracy for movement decoding (A) and identity decoding (B) during the retention period at p < 0.05, FWE corrected. Only clusters encompassing at least 50 voxels are displayed.

| (A) Movement decoding |  | Peak MNI coordinates |  |  |  |
| --- | --- | --- | --- | --- | --- |
| Cluster size | Anatomical region | *X* | *Y* | *Z* | *z*-score |
| 19015 | Left Middle Temporal Gyrus | -48 | -60 | 4 | 7.82 |
|  | Left Superior Parietal Lobule | -36 | -46 | 56 | 7.77 |
|  | Right Inferior Occipital Gyrus | 50 | -70 | 6 | 7.19 |
| 5587 | Left Precentral Gyrus | -34 | -8 | 58 | 7.43 |
|  | Right Precentral Gyrus | 30 | -2 | 54 | 7.11 |
|  | Left Supplementary Motor Cortex | -2 | 2 | 58 | 6.81 |
| 135 | Right Temporal Pole | 20 | 20 | -38 | 6.40 |
| 305 | Left Inferior Occipital Gyrus | -30 | -94 | -22 | 5.54 |
| 313 | R Inferior Frontal Gyrus / white matter | 36 | 16 | 20 | 5.03 |
| 185 | Left posterior cingulate gyrus | -8 | -42 | -36 | 4.86 |
| 293 | Right supramarginal gyrus | 64 | -28 | 36 | 4.79 |
| 61 | Right Exterior Cerebellum | 36 | -62 | -56 | 4.61 |
| (B) Identity decoding |  | Peak MNI coordinates |  |  |  |
| Cluster size | Anatomical region | *X* | *Y* | *Z* | *z*-score |
| 1170 | Left Middle Frontal Gyrus | -40 | 4 | 40 | 7.46 |
|  | Left Inferior Frontal Gyrus, pars opercularis | -54 | 24 | 20 | 5.33 |
| 68 | Left Supplementary Motor Area | -2 | 14 | 54 | 5.21 |

**Figure 2.**
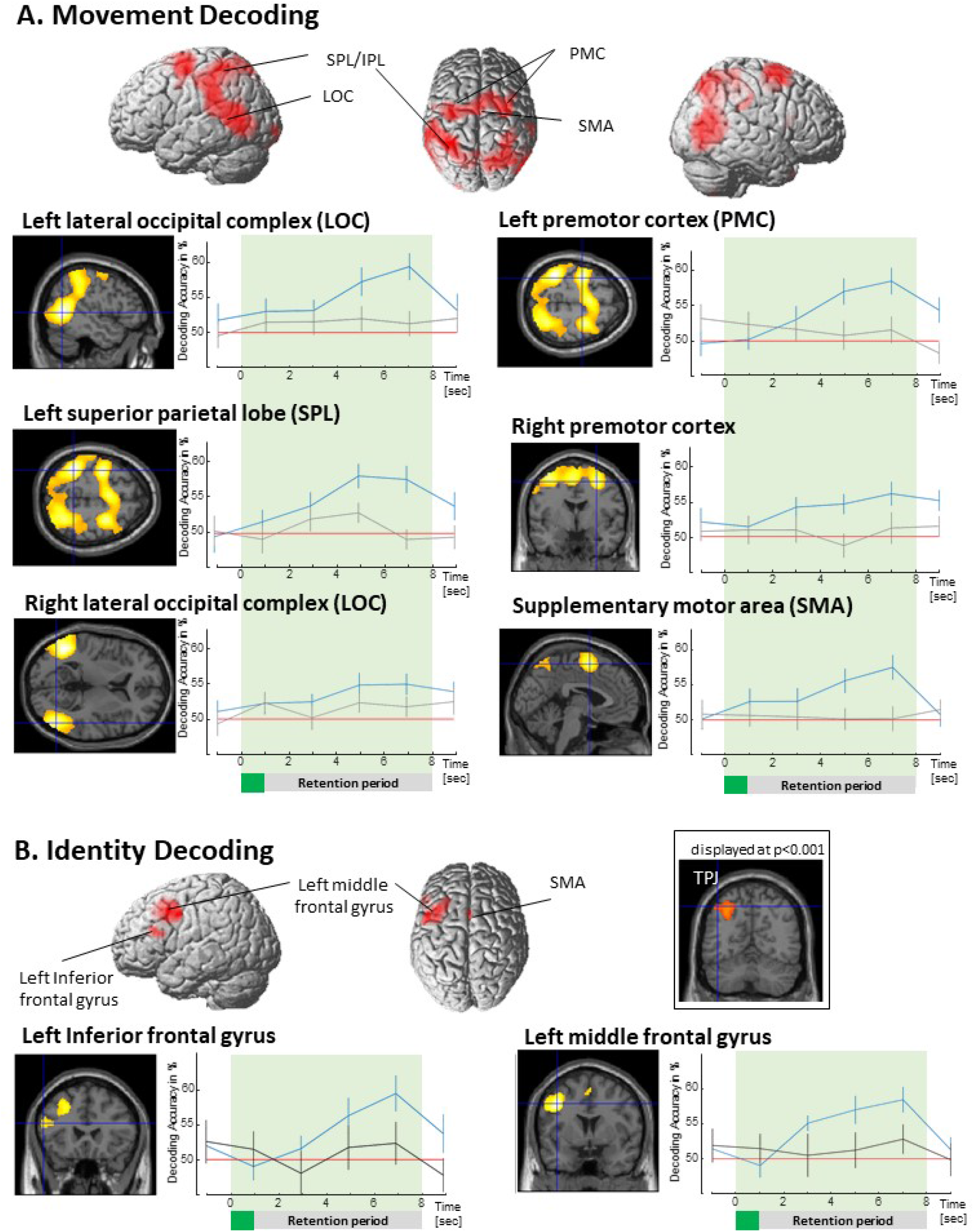
**A**. Brain areas representing information about the type of movement held in working memory. Blue lines represent decoding accuracy from peak voxels for memorized movement. Grey lines represent decoding accuracy from the same areas for non-memorized movement. **B**. Brain areas representing information about whether the movement held in working memory is associated with the self vs. others. Blue lines represent decoding accuracy from peak voxels between self and others as a factor of time. Black lines represent decoding accuracy from the same areas between two control (“other”) identities: Pat and Doc. All results displayed at p < 0.05, FWE corrected, error bars indicate SEM.

Time courses of decoding accuracies for the identified regions are displayed in Figure 2. The delayed build-up of decoding accuracies along with the levelling off until the end of the retention period indicate that the decoding results are specific to a WM representation and do not stem from mere perceptual processing (see also control analyses).

### Identity Decoding

In order to localize brain areas representing identity-related information, we computed two decoding analyses: (1) Self versus Other identities, and (2) Pat versus Doc. Brain regions representing self-related information should show above-chance level decoding accuracies only in (1), where activation patterns of a self-related condition are compared to activation pattern elicited by Others. In turn, regions coding self-related information are assumed to not show above-chance decoding accuracies when activation pattern of two Other persons are compared. Thereby regions revealed in (1) and not in (2) code not only the difference between two identities, but the difference between movements encoded for oneself and movements encoded for other persons. A corresponding t-contrast of decoding accuracies against chance-level across the WM retention period (p<0.05, FWE corrected, min. cluster size = 50) revealed two clusters for self vs. others: One cluster spanning the left middle frontal gyrus and the left inferior frontal gyrus, pars opercularis, and a smaller cluster in the left supplementary motor area (**Figure 2, Table 1**).

Following our a priori hypothesis we inspected the data also at a lower threshold to check for above change decoding accuracy in the TPJ and vmPFC, which might have been missed due to conservative FWE correction threshold. Indeed, this revealed a cluster in the TPJ, more specifically angular gyrus (p<0.001, FWE-corrected on the cluster level; peak coordinates: x=-30, y=-64, z=48, size = 643 voxel, z = 436). No above chance decoding was found in the vmPFC.

Testing for above-chance decoding between the two Other-identities Pat vs. Doc yielded no significant clusters. This suggests that the regions revealed in the Self versus Other decoding are specific to the coding of self-related information.

### Control analyses: Decoding of non-memorized content

We ran three control decoding analyses to test for the specificity of the main results: the (1) Non-memorized Movement Decoding, (2) Non-memorized Identity Decoding: self vs. others, and (3) Non-memorized Identity Decoding: Pat vs. Doc. These analyses contained the same amount of data and the same processing steps as the three main analyses but tested if information can be decoded on the corresponding non-memorized stimuli. No significant clusters of activation were found in either of the analyses.

### Univariate analysis

Testing in an assumption-free manner throughout the whole brain for the self > other contrast of the retro-cue revealed activation clusters in the left vmPFC (x=-6, y=58, z=4, cluster size = 113 voxel, z-score = 3.88; **Figure 3**), the left superior frontal gyrus (x=-20, y=34, z=42, cluster size = 231 voxel, z-score = 4.44) and in the right superior frontal gyrus (cluster1: x=22, y=42, z=44, cluster size = 150 voxel, z-score = 4.56), at p<0.05 FWE-corrected on the cluster level (clusters did not survive *p*<0.05 FWE correction for the whole brain). Contrasting self vs. other for the WM retention period did not reveal any significant clusters.

**Figure 3.**
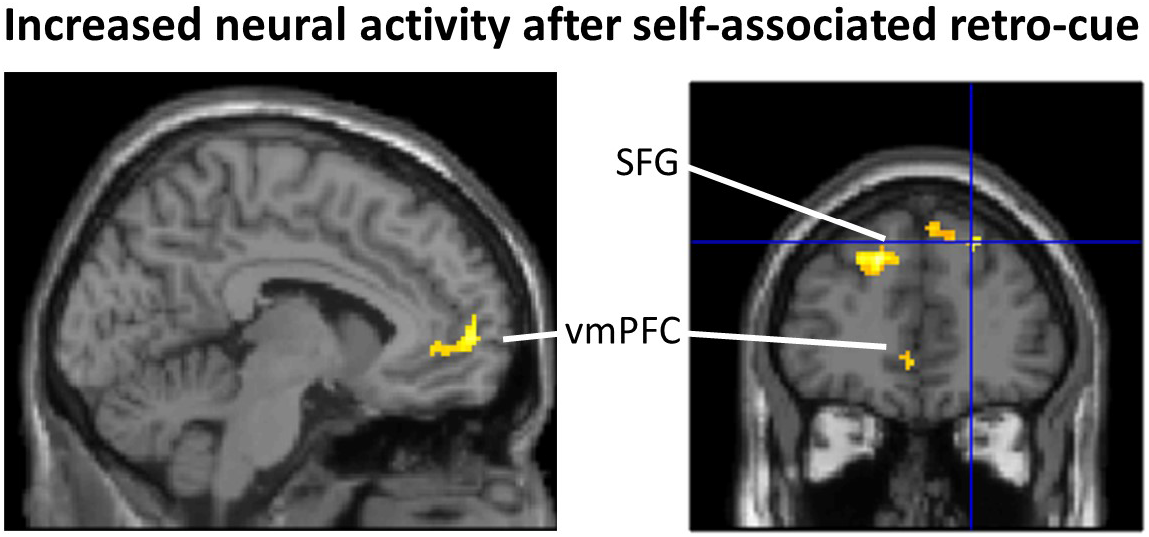
Presentation of a self-associated retro-cue led to increased activity in the ventral medial prefrontal cortex and bilaterally in the superior frontal gyri. p<0.05 FWE-corrected on the cluster level.

Testing for differences between the two stranger-identity conditions (Pat and Doc, in both directions) did not reveal any significant clusters neither for the retro-cue, nor throughout the working memory retention period (p<0.001 uncorrected).

## Discussion

We conducted an fMRI study and used MVPA to test which brain areas code information about whether a memorized movement was represented in WM as performed by oneself or by someone else. First, we tested for the brain network in which activation patterns relate to the retention of specific movements (e.g. chopping, painting, etc.: Movement Decoding). We found a network of regions, related to higher-order perceptual and multimodal integration processing of movements, such as the superior temporal sulcus, middle temporal cortex, extrastriate body area, intraparietal sulcus, as well as supplementary and premotor/motor cortices. Testing for brain regions that exhibit distinct activations for movements associated with oneself as compared to those associated with another identity (Identity Decoding), our analyses revealed the left middle frontal gyrus, inferior frontal gyrus, and supplementary motor area. Finally, we found increased activation in the vmPFC in response to the presentation of a self-associated retro-cue, as well as in bilateral SFG, however not during the WM retention period.

### Movement Decoding

Multiple WM MVPA studies indicate that the brain regions that exhibit activity related to specific WM contents partly overlap with the brain regions involved in the perceptual processing of corresponding stimulus material (Lee & Baker, 2016). It has been suggested that the amount of overlap is determined by the degree of abstractness in which contents are retained, e.g. in a rather sensory format, like imagining the visual appearance of a shape, or in a rather symbolic or language format, like used when rehearsing a telephone number (Christophel, Klink, Spitzer, Roelfsema, & Haynes, 2017). Our movement decoding analysis extends this literature, by testing what brain regions exhibit codes for the retention of more complex and more naturalistic material, namely specific movements. Indeed, the regions revealed by our analysis partly overlap with regions known for perceptual processing of biological motion and action observation (Caspers, Zilles, Laird, & Eickhoff, 2010; E. Grossman et al., 2000; E. D. Grossman & Blake, 2002; Lu et al., 2016; Pelphrey, Morris, Michelich, Allison, & McCarthy, 2005; Saygin, 2007a; Saygin, Wilson, Hagler, Bates, & Sereno, 2004; Vaina, Solomon, Chowdhury, Sinha, & Belliveau, 2001; van Kemenade, Muggleton, Walsh, & Saygin, 2012).

Finding only a small cluster in early visual areas indicates that the mere visual appearance of the stimulus material plays a minor role for the given WM representations, in contrast to the retention of static visual stimulus material (e.g. Albers, Kok, Toni, Dijkerman, & De Lange, 2013; Christophel et al., 2012). As expected, our analysis revealed bilateral clusters in the lateral occipital cortex (LOC), including the motion-sensitive human middle temporal (hMT) region and the extrastriate body area (EBA), which have been shown to be involved in the visual processing of human body parts (Limanowski & Blankenburg, 2016). The identified cluster extends to the posterior superior temporal sulcus (pSTS), which has been found to be sensitive to movements and goal-directed actions (Pelphrey et al., 2005; Pitcher & Ungerleider, 2020; Saygin, 2007b; Shultz, Lee, Pelphrey, & McCarthy, 2011; Vaina et al., 2001). Superior- and intraparietal lobe regions (SPL/IPL) have been found in visual and tactile WM tasks where spatial information was retained, and are well known for their involvement in the remapping of spatial coordinate systems between allocentric and egocentric perspectives, particularly important during movements (Christophel et al., 2012; Heed, Buchholz, Engel, & Röder, 2015; Schmidt & Blankenburg, 2018). Finally, as expected, our analysis revealed pre- and supplementary motor areas. These regions have been found to contribute to different types of WM representations (Schmidt et al., 2017; Uluç, Schmidt, Wu, & Blankenburg, 2018), where they are thought to reflect attentional top-down mechanisms of mental rehearsal processes (Compare: Schmidt, Schröder, Reinhardt, & Blankenburg, 2020). These regions are also well known for their contributions to action planning and motor coordination. Therefore, they most likely reflect the mental simulation of the memorized movement (Rizzolatti & Sinigaglia, 2010, but see also: Cook, Bird, Catmur, Press, & Heyes, 2014).

Taken together, the identified network of regions partly overlaps with regions involved in perceptual processing of dot-motion movement-stimuli (Lu et al., 2016) and regions involved in mental simulation of movements.

### Identity Decoding

The main goal of our study was to test which brain areas’ activity allows us to distinguish whether a memorized full-body movement was represented as a movement performed by oneself or performed by someone else. Our results demonstrate that even an arbitrary movement, if it is interpreted as one’s own movement, induces different brain activation than a movement not associated with oneself. We found above-chance decoding accuracy in one cluster spanning the left middle frontal gyrus and the left inferior frontal gyrus pars opercularis, and a second cluster in the left supplementary motor area. All of these areas (especially the IFG pars opercularis) have previously been identified to be active during action observation and imitation (Caspers et al., 2010; Johnson-Frey et al., 2003; Molnar-Szakacs, Iacoboni, Koski, & Mazziotta, 2005). Furthermore, there is evidence that IFG pars opercularis and MFG constitute parts of the human mirror neuron system, as indicated by ALE-meta-analysis by Molenberghs et al. (2012). The differences in IFG, MFG, and SMA were revealed by our MVPA while the corresponding univariate comparison did not reveal differences. This shows that our results do not simply reflect that the activity was greater in these areas, when self-than other-associated movements were retained in WM. Instead, they suggest that neural codes, or the activation pattern within these areas, differ when one observes a movement that is represented as “myself” versus “other”, which can be detected by the higher sensitivity of MVPA as compared to univariate analyses.

All movements used in our study were displayed from the third person (allocentric) perspective, that is, how we usually perceive other people. A situations in which we observe our bodies acting from that perspective is when we see ourselves in a mirror or on a video. However, in these two kinds of situations, we observe ourselves performing movements that we already know from experience – i.e. we performed them when they were recorded (in case of a recently recorded video), or that we’re performing right now (in case of a mirror). It has been found that activity of the mirror neurons network, including the IFG, is increased when observing actions that are familiar to us (Calvo-Merino, Glaser, Grèzes, Passingham, & Haggard, 2004; Calvo-Merino, Grèzes, Glaser, Passingham, & Haggard, 2006; Cross, Hamilton, & Grafton, 2006; Cross, Hamilton, Kraemer, Kelley, & Grafton, 2009). However, in our study, no such effect could directly account for the results, because our design excluded the effect of familiarity of the actions associated with each identity, because for every participant the same actions were equally often associated with all identities across trials. Therefore, our findings in the IFG do not simply reflect motor familiarity, instead they reflect in a well-controlled manner whether a seen action is represented as self- or other-related.

The second region we expected to be modulated by self-association was the TPJ. With a more liberal significance threshold we found that activity in the left angular gyrus, which constitutes a dorsal posterior part of the TPJ, differentiated between maintaining representations of self- and other-associated movements. Bilateral TPJ has been suggested as an area that is responsible for implementing domain-general self-other distinction, as shown by a large variety of tasks involving self-body perception, self-agency, and mindreading (Quesque & Brass, 2019). Some research points to the role of the left TPJ in third-person self-representation (Ganesh et al., 2011), but more research is needed to fully establish this link. Previous studies on perception of actions performed by self or others that have identified TPJ activity mainly used videos and animations of participants’ real movements – stimuli that were portraying highly familiar movement patterns (e.g. Bischoff et al., 2012; Macuga & Frey, 2011; Sugiura et al., 2006). These studies typically argued that the role of TPJ in self-other distinction relies on detecting motor familiarity, and specifically on comparing predicted and observed body movements. However, because all animations used in our study reflected kinematics that were equally unfamiliar to participants (none of them was based on their real kinematics), our results support the view that the TPJ’s role in representing the distinction between self and others is more domain-general as advocated by (Quesque & Brass, 2019), and is not limited to a mechanisms that rely on familiarity of observed movements.

Surprisingly, in behavioral data we found that participants performed slightly worse in the forced-choice task at the end of the WM delay period, when memorizing self-associated movements than when they were memorizing other-associated movements. While this finding promotes the view that self-associated movements are distinctly processed, previous work has indicated that self-related material is processed in a priviledged way and one could therefore expect a higher performance for WM of self-related material (Sui & Humphreys, 2015; Symons & Johnson, 1997). One interpretation that could explain this effect would be that the unfamiliar movements that had to be actively associated with oneself might cause interference with knowledge about one’s real body dynamics. Therefore the WM representation might have been influenced by one’s own sensorimotor body representation (De Vignemont, 2010, 2018; Dewey & Knoblich, 2016; Gallagher, 2005; Kanayama & Hiromitsu, 2021; Schwoebel & Coslett, 2005), which would make it more difficult to match the memorized movement with the comparison stimulus. This difference in performance should not have affected the main neuroimaging results, as a control analysis modelling only the correct trials confirmed above chance decoding in the three reported regions.

We did not find distinctive representations of self during the WM retention period in one area in which we expected to find them: in vmPFC. Previous studies found that aspects of the conceptual self (self-related adjectives, autobiographical memories, etc.) are usually associated with increased activity in midline structures (Qin & Northoff, 2011; Qin et al., 2020): precuneus and vmPFC, the latter of which is often regarded as the central node for self-related processing (for a discussion see: D’Argembeau, 2013; Martinelli et al., 2013; Northoff et al., 2006). However, studies which investigated aspects of the self related to one’s body, rather than personality, rarely find modulation of activity in the vmPFC or other midline structures (Gillihan & Farah, 2005; Hu et al., 2016; Legrand & Ruby, 2009; Qin et al., 2020). A possible interpretation of these results is that midline structures, in particular vmPFC, are more relevant for abstract and conceptual (e.g., semantic) information about the self than for the less abstract and more naturalistic bodily information associated with the self used in the study at hand. However, in univariate analyses we did find increased activation in the vmPFC following presentation of a self-associated retro-cue. In our study, the retro-cue displayed one of the colors that prior to the experiment were associated with the self or the other identities. As such, it served the function of an abstract symbol referring to the self or others, in the same way as geometrical shapes that have been used in previous fMRI research on processing of arbitrary self-associated stimuli (self-prioritization effect: Sui et al., 2013; Yankouskaya et al., 2017, see also: Lockwood et al., 2018; Yin et al., 2021). Moreover, the specific vmPFC cluster detected in our study overlapped with the cluster found in previous studies where processing of self-associated shapes was tested (Sui et al., 2013). This pattern of results supports previous findings that vmPFC is primarily associated with self-related processing of conceptual aspects of the self, and not necessarily bodily aspects. Moreover, the finding that self-association is related to increased activation in vmPFC after presentation of the retro-cue, but not throughout the whole retention period, suggests that vmPFC is involved in selecting self-associations for retention in working memory (Myers, Stokes, & Nobre, 2017), but not in maintaining them. This interpretation is consistent with previous research on the influence of self-relatedness on memory, which indicates that vmPFC is involved in encoding and retrieval, but not maintenance of self-related information (Kalenzaga et al., 2015; Leshikar & Duarte, 2012; Philippi, Duff, Denburg, Tranel, & Rudrauf, 2012; Yaoi, Osaka, & Osaka, 2015).

## Conclusions

We conducted a study in which we tested which brain areas display different coding depending on whether a memorized action (seen from the third-person perspective) is represented as being performed by oneself or by someone else. Crucially and in contrast to previous studies investigating this topic, as the same movements were associated with the self and with other identities in different trials, our study design rendered confounding influence of motor and perceptual familiarity unlikely. Our findings indicate that self-associated movements are differentially represented during the working memory retention period through left hemispheric activation patterns in frontal areas responsible for action recognition and motor mirroring (MFG, IFG and SMA). We further found increased activity in the vmPFC directly after the presentation of the retro-cue that instructed to recall a self-associated movement but no difference across the working memory retention period. This suggests that the influence of the vmPFC in our task might be limited to selecting a self-associated movement for working memory, and not for maintaining it in working memory.

## Data and Code Availability Statement

The raw data generated during the current study are not publicly available due to data protection laws, but normalized data and the used scripts are available from the corresponding author on request.

## Acknowledgments

This research was funded by the Australian Research Council grant DP160102770.

## References

Albers, A. M., Kok, P., Toni, I., Dijkerman, H. C., & De Lange, F. P. (2013). Shared representations for working memory and mental imagery in early visual cortex. Current Biology, 23(15), 1427–1431.

Baumeister, R. F. (1999). The self in social psychology. Philadelphia: Psychology Press.

Bischoff, M., Zentgraf, K., Lorey, B., Pilgramm, S., Balser, N., Baumgartner, E., … Munzert, J. (2012). Motor familiarity: Brain activation when watching kinematic displays of one’s own movements. Neuropsychologia, 50(8), 2085–2092.

Blanke, O. (2012). Multisensory brain mechanisms of bodily self-consciousness. Nat Rev Neurosci, 13(8), 556–571. doi:10.1038/nrn3292

Blanke, O., Slater, M., & Serino, A. (2015). Behavioral, Neural, and Computational Principles of Bodily Self-Consciousness. Neuron, 88(1), 145–166. doi:10.1016/j.neuron.2015.09.029

Calvo-Merino, B., Glaser, D. E., Grèzes, J., Passingham, R. E., & Haggard, P. (2004). Action observation and acquired motor skills: an FMRI study with expert dancers. Cerebral cortex, 15(8), 1243–1249.

Calvo-Merino, B., Grèzes, J., Glaser, D. E., Passingham, R. E., & Haggard, P. (2006). Seeing or doing? Influence of visual and motor familiarity in action observation. Current Biology, 16(19), 1905–1910.

Caspers, S., Zilles, K., Laird, A. R., & Eickhoff, S. B. (2010). ALE meta-analysis of action observation and imitation in the human brain. Neuroimage, 50(3), 1148–1167.

Chang, C.-C., & Lin, C.-J. (2011). LIBSVM: A library for support vector machines. ACM transactions on intelligent systems and technology (TIST), 2(3), 27.

Christoff, K., Cosmelli, D., Legrand, D., & Thompson, E. (2011). Specifying the self for cognitive neuroscience. Trends Cogn Sci, 15(3), 104–112. doi:10.1016/j.tics.2011.01.001

Christophel, T. B., Hebart, M. N., & Haynes, J.-D. (2012). Decoding the contents of visual short-term memory from human visual and parietal cortex. Journal of Neuroscience, 32(38), 12983–12989.

Christophel, T. B., Klink, P. C., Spitzer, B., Roelfsema, P. R., & Haynes, J.-D. (2017). The distributed nature of working memory. Trends in Cognitive Sciences, 21(2), 111–124.

Cook, R., Bird, G., Catmur, C., Press, C., & Heyes, C. (2014). Mirror neurons: from origin to function. Behavioral and brain sciences, 37(2), 177–192.

Cross, E. S., Hamilton, A. F. d. C., & Grafton, S. T. (2006). Building a motor simulation de novo: observation of dance by dancers. Neuroimage, 31(3), 1257–1267.

Cross, E. S., Hamilton, A. F. d. C., Kraemer, D. J., Kelley, W. M., & Grafton, S. T. (2009). Dissociable substrates for body motion and physical experience in the human action observation network. European journal of neuroscience, 30(7), 1383–1392.

D’Argembeau, A. (2013). On the role of the ventromedial prefrontal cortex in self-processing: the valuation hypothesis. Frontiers in human neuroscience, 7, 372.

De Vignemont, F. (2010). Body schema and body image—Pros and cons. Neuropsychologia, 48(3), 669–680.

De Vignemont, F. (2018). Mind the body: An exploration of bodily self-awareness: Oxford University Press.

Dewey, J. A., & Knoblich, G. (2016). 17 Representation of Self versus Others’ Actions. Shared Representations: Sensorimotor Foundations of Social Life, 351.

Gallagher, S. (2000). Philosophical conceptions of the self: implications for cognitive science. Trends Cogn Sci, 4(1), 14–21. Retrieved from https://www.ncbi.nlm.nih.gov/pubmed/10637618

Gallagher, S. (2005). How the body shapes the mind. Oxford; New York: Clarendon Press.

Ganesh, S., van Schie, H. T., de Lange, F. P., Thompson, E., & Wigboldus, D. H. (2011). How the human brain goes virtual: Distinct cortical regions of the person-processing network are involved in self-identification with virtual agents. Cerebral cortex, 22(7), 1577–1585.

Gillihan, S. J., & Farah, M. J. (2005). Is self special? A critical review of evidence from experimental psychology and cognitive neuroscience. Psychological Bulletin, 131(1), 76.

Grossman, E., Donnelly, M., Price, R., Pickens, D., Morgan, V., Neighbor, G., & Blake, R. (2000). Brain areas involved in perception of biological motion. Journal of Cognitive Neuroscience, 12(5), 711–720.

Grossman, E. D., & Blake, R. (2002). Brain areas active during visual perception of biological motion. Neuron, 35(6), 1167–1175.

Hebart, M. N., Görgen, K., & Haynes, J.-D. (2015). The Decoding Toolbox (TDT): a versatile software package for multivariate analyses of functional imaging data. Frontiers in neuroinformatics, 8, 88.

Heed, T., Buchholz, V. N., Engel, A. K., & Röder, B. (2015). Tactile remapping: from coordinate transformation to integration in sensorimotor processing. Trends in Cognitive Sciences, 19(5), 251–258.

Hu, C., Di, X., Eickhoff, S. B., Zhang, M., Peng, K., Guo, H., & Sui, J. (2016). Distinct and common aspects of physical and psychological self-representation in the brain: A meta-analysis of self-bias in facial and self-referential judgements. Neuroscience & biobehavioral reviews, 61, 197–207.

Johnson-Frey, S. H., Maloof, F. R., Newman-Norlund, R., Farrer, C., Inati, S., & Grafton, S. T. (2003). Actions or hand-object interactions? Human inferior frontal cortex and action observation. Neuron, 39(6), 1053–1058.

Kalenzaga, S., Sperduti, M., Anssens, A., Martinelli, P., Devauchelle, A.-D., Gallarda, T., … Krebs, M.-O. (2015). Episodic memory and self-reference via semantic autobiographical memory: insights from an fMRI study in younger and older adults. Frontiers in behavioral neuroscience, 8, 449.

Kanayama, N., & Hiromitsu, K. (2021). Triadic body representations in the human cerebral cortex and peripheral nerves. Body Schema and Body Image: New Directions, 133.

Kriegeskorte, N., Goebel, R., & Bandettini, P. (2006). Information-based functional brain mapping. Proceedings of the National Academy of Sciences, 103(10), 3863–3868.

Lee, S.-H., & Baker, C. I. (2016). Multi-voxel decoding and the topography of maintained information during visual working memory. Frontiers in systems neuroscience, 10, 2.

Legrand, D., & Ruby, P. (2009). What is self-specific? Theoretical investigation and critical review of neuroimaging results. Psychological review, 116(1), 252.

Leshikar, E. D., & Duarte, A. (2012). Medial prefrontal cortex supports source memory accuracy for self-referenced items. Social neuroscience, 7(2), 126–145.

Limanowski, J., & Blankenburg, F. (2013). Minimal self-models and the free energy principle. Front Hum Neurosci, 7, 547. doi:10.3389/fnhum.2013.00547

Limanowski, J., & Blankenburg, F. (2016). Integration of visual and proprioceptive limb position information in human posterior parietal, premotor, and extrastriate cortex. Journal of Neuroscience, 36(9), 2582–2589.

Lockwood, P. L., Wittmann, M. K., Apps, M. A., Klein-Flügge, M. C., Crockett, M. J., Humphreys, G. W., & Rushworth, M. F. (2018). Neural mechanisms for learning self and other ownership. Nature communications, 9(1), 1–11.

Lu, X., Huang, J., Yi, Y., Shen, M., Weng, X., & Gao, Z. (2016). Holding biological motion in working memory: An fMRI study. Frontiers in human neuroscience, 10, 251.

Macey, P. M., Macey, K. E., Kumar, R., & Harper, R. M. (2004). A method for removal of global effects from fMRI time series. Neuroimage, 22(1), 360–366.

Macuga, K. L., & Frey, S. H. (2011). Selective responses in right inferior frontal and supramarginal gyri differentiate between observed movements of oneself vs. another. Neuropsychologia, 49(5), 1202–1207.

Martinelli, P., Sperduti, M., & Piolino, P. (2013). Neural substrates of the self-memory system: new insights from a meta-analysis. Hum Brain Mapp, 34(7), 1515–1529. doi:10.1002/hbm.22008

Mattan, B., Quinn, K. A., Apperly, I. A., Sui, J., & Rotshtein, P. (2015). Is it always me first? Effects of self-tagging on third-person perspective-taking. J Exp Psychol Learn Mem Cogn, 41(4), 1100–1117. doi:10.1037/xlm0000078

Molenberghs, P., Cunnington, R., & Mattingley, J. B. (2012). Brain regions with mirror properties: a meta-analysis of 125 human fMRI studies. Neuroscience & biobehavioral reviews, 36(1), 341–349.

Molnar-Szakacs, I., Iacoboni, M., Koski, L., & Mazziotta, J. C. (2005). Functional segregation within pars opercularis of the inferior frontal gyrus: evidence from fMRI studies of imitation and action observation. Cerebral cortex, 15(7), 986–994.

Myers, N. E., Stokes, M. G., & Nobre, A. C. (2017). Prioritizing information during working memory: beyond sustained internal attention. Trends in Cognitive Sciences, 21(6), 449–461.

Neisser, U. (1988). Five kinds of self-knowledge. Philosophical Psychology, 1(1), 35–59.

Northoff, G., Heinzel, A., De Greck, M., Bermpohl, F., Dobrowolny, H., & Panksepp, J. (2006). Self-referential processing in our brain—a meta-analysis of imaging studies on the self. Neuroimage, 31(1), 440–457.

Payne, S., Tsakiris, M., & Maister, L. (2017). Can the self become another? Investigating the effects of self-association with a new facial identity. Q J Exp Psychol (Hove), 70(6), 1085–1097. doi:10.1080/17470218.2015.1137329

Pelphrey, K. A., Morris, J. P., Michelich, C. R., Allison, T., & McCarthy, G. (2005). Functional anatomy of biological motion perception in posterior temporal cortex: an fMRI study of eye, mouth and hand movements. Cerebral cortex, 15(12), 1866–1876.

Philippi, C. L., Duff, M. C., Denburg, N. L., Tranel, D., & Rudrauf, D. (2012). Medial PFC damage abolishes the self-reference effect. Journal of Cognitive Neuroscience, 24(2), 475–481.

Pitcher, D., & Ungerleider, L. G. (2020). Evidence for a Third Visual Pathway Specialized for Social Perception. Trends in Cognitive Sciences.

Qin, P., & Northoff, G. (2011). How is our self related to midline regions and the default-mode network? Neuroimage, 57(3), 1221–1233.

Qin, P., Wang, M., & Northoff, G. (2020). Linking Bodily, Environmental and Mental States In the Self–A Three-level Model Based on A Meta-analysis. Neuroscience & biobehavioral reviews.

Quesque, F., & Brass, M. (2019). The role of the temporoparietal junction in self-other distinction. Brain topography, 32(6), 943–955.

Rizzolatti, G., & Sinigaglia, C. (2010). The functional role of the parieto-frontal mirror circuit: interpretations and misinterpretations. Nature Reviews Neuroscience, 11(4), 264–274.

Saygin, A. P. (2007a). Brain areas involved in biological motion perception: What is involved and what is necessary. Journal of vision, 7(9), 492–492.

Saygin, A. P. (2007b). Superior temporal and premotor brain areas necessary for biological motion perception. Brain, 130(9), 2452–2461.

Saygin, A. P., Wilson, S. M., Hagler, D. J., Bates, E., & Sereno, M. I. (2004). Point-light biological motion perception activates human premotor cortex. Journal of Neuroscience, 24(27), 6181–6188.

Schäfer, S., Wentura, D., & Frings, C. (2017). Distinctiveness effects in self-prioritization. Visual Cognition, 25(1-3), 399–411.

Schmidt, T. T., & Blankenburg, F. (2018). Brain regions that retain the spatial layout of tactile stimuli during working memory–A ‘tactospatial sketchpad’? Neuroimage.

Schmidt, T. T., Schröder, P., Reinhardt, P., & Blankenburg, F. (2020). Rehearsal of tactile working memory: Premotor cortex recruits two dissociable neuronal content representations. Human brain mapping.

Schmidt, T. T., Wu, Y.-h., & Blankenburg, F. (2017). Content-specific codes of parametric vibrotactile working memory in humans. Journal of Neuroscience, 1167–1117.

Schwoebel, J., & Coslett, H. B. (2005). Evidence for multiple, distinct representations of the human body. Journal of Cognitive Neuroscience, 17(4), 543–553.

Shi, Z., Zhou, A., Han, W., & Liu, P. (2011). Effects of ownership expressed by the first-person possessive pronoun. Conscious Cogn, 20(3), 951–955. doi:10.1016/j.concog.2010.12.008

Shultz, S., Lee, S. M., Pelphrey, K., & McCarthy, G. (2011). The posterior superior temporal sulcus is sensitive to the outcome of human and non-human goal-directed actions. Social Cognitive and Affective Neuroscience, 6(5), 602–611.

Sugiura, M., Sassa, Y., Jeong, H., Miura, N., Akitsuki, Y., Horie, K., … Kawashima, R. (2006). Multiple brain networks for visual self-recognition with different sensitivity for motion and body part. Neuroimage, 32(4), 1905–1917.

Sui, J., He, X., & Humphreys, G. W. (2012). Perceptual effects of social salience: evidence from self-prioritization effects on perceptual matching. J Exp Psychol Hum Percept Perform, 38(5), 1105–1117. doi:10.1037/a0029792

Sui, J., & Humphreys, G. W. (2015). The Integrative Self: How Self-Reference Integrates Perception and Memory. Trends Cogn Sci, 19(12), 719–728. doi:10.1016/j.tics.2015.08.015

Sui, J., Liu, M., Mevorach, C., & Humphreys, G. W. (2015). The salient self: The left intraparietal sulcus responds to social as well as perceptual-salience after self-association. Cerebral cortex, 25(4), 1060–1068.

Sui, J., Rotshtein, P., & Humphreys, G. W. (2013). Coupling social attention to the self forms a network for personal significance. Proc Natl Acad Sci U S A, 110(19), 7607–7612. doi:10.1073/pnas.1221862110

Symons, C. S., & Johnson, B. T. (1997). The self-reference effect in memory: a meta-analysis. Psychol Bull, 121(3), 371–394. Retrieved from https://www.ncbi.nlm.nih.gov/pubmed/9136641

Tsakiris, M. (2010). My body in the brain: a neurocognitive model of body-ownership. Neuropsychologia, 48(3), 703–712.

Tsakiris, M. (2017). The multisensory basis of the self: from body to identity to others. The Quarterly Journal of Experimental Psychology, 70(4), 597–609.

Uluç, I., Schmidt, T. T., Wu, Y.-h., & Blankenburg, F. (2018). Content-specific codes of parametric auditory working memory in humans. Neuroimage, 183, 254–262.

Vaina, L. M., Solomon, J., Chowdhury, S., Sinha, P., & Belliveau, J. W. (2001). Functional neuroanatomy of biological motion perception in humans. Proceedings of the National Academy of Sciences, 98(20), 11656–11661.

van Kemenade, B. M., Muggleton, N., Walsh, V., & Saygin, A. P. (2012). Effects of TMS over premotor and superior temporal cortices on biological motion perception. Journal of Cognitive Neuroscience, 24(4), 896–904.

Vanrie, J., & Verfaillie, K. (2004). Perception of biological motion: A stimulus set of human point-light actions. Behavior Research Methods, Instruments, & Computers, 36(4), 625–629.

Woźniak, M., & Hohwy, J. (2020). Stranger to my face: top-down and bottom-up effects underlying prioritization of images of one’s face PLoS One.

Woźniak, M., & Knoblich, G. (2019). Self-prioritization of fully unfamiliar stimuli. Quarterly Journal of Experimental Psychology. doi:https://doi.org/10.1177/1747021819832981

Woźniak, M., Kourtis, D., & Knoblich, G. (2018). Prioritization of arbitrary faces associated to self: An EEG study. PLoS One, 13(1), e0190679.

Yankouskaya, A., Humphreys, G., Stolte, M., Stokes, M., Moradi, Z., & Sui, J. (2017). An anterior–posterior axis within the ventromedial prefrontal cortex separates self and reward. Social Cognitive and Affective Neuroscience, 12(12), 1859–1868.

Yankouskaya, A., & Sui, J. (2021). Self-Positivity or Self-Negativity as a Function of the Medial Prefrontal Cortex. Brain Sciences, 11(2), 264.

Yaoi, K., Osaka, M., & Osaka, N. (2015). Neural correlates of the self-reference effect: evidence from evaluation and recognition processes. Frontiers in human neuroscience, 9, 383.

Yin, S., Bi, T., Chen, A., & Egner, T. (2021). Ventromedial prefrontal cortex drives the prioritization of self-associated stimuli in working memory. Journal of Neuroscience, 41(9), 2012–2023.

Zhao, S., Uono, S., Li, C., Yoshimura, S., & Toichi, M. (2018). The influence of self-referential processing on attentional orienting in frontoparietal networks. Frontiers in human neuroscience, 12.

Zhou, A., Shi, Z., Zhang, P., Liu, P., Han, W., Wu, H., … Xia, R. (2010). An ERP study on the effect of self-relevant possessive pronoun. Neurosci Lett, 480(2), 162–166. doi:10.1016/j.neulet.2010.06.033

